# Learning the mechanism of collective microbial function via random community-media pairing

**DOI:** 10.1101/2025.11.25.690499

**Authors:** Luoqi Wang, Seppe Kuehn, Mikhail Tikhonov

**Affiliations:** Department of Physics, Washington University in St. Louis, St. Louis, MO 63130, USA; Department of Ecology and Evolution, The University of Chicago, Chicago, IL 60637, USA; Center for the Physics of Evolving Systems, The University of Chicago, Chicago, IL 60637, USA; Center for Living Systems, The University of Chicago, Chicago, IL 60637, USA; National Institute for Theory and Mathematics in Biology, Northwestern University and The University of Chicago, Chicago, IL 60611, USA

## Abstract

In microbial systems, many biochemical functions arise from pathways encoded and executed at the community level. The collective nature of these functions complicates bottom-up efforts to determine each species’ contribution. Previous work has shown that regression over randomly sampled datasets of collective functions succeeds at predicting those functions. Building on this top-down idea, this paper asks whether regression can also reveal mechanistic insight into a cross-feeding relationship. For this, we propose extending the random sampling method to vary the growth environment, cultivating each community in the spent medium of another randomly constructed community. With a model-based analysis, we show that the new protocol extracts more mechanistic information, enabling assignment of species to the correct cross-feeding pathway steps and identification of species essential to the collective function, both achieved with simple LASSO regressions. More generally, our work illustrates that the utility of machine learning-based approaches can be greatly enhanced by a synergistic experimental design.

## I. INTRODUCTION

Microbial communities, particularly those assembled naturally, are often very complex. The complex interactions among microbial strains often lead to collective functions that are absent when examining individual strains but emerge at community level. Examples of such collective functions include toxin production, toxin degradation, or invasion resistance. Surprisingly, despite the higher-order interactions in the communities, these collective functions can sometimes be efficiently predicted with methods as simple as regression over presence-absence information of randomly-sampled species subsets [1] (FIG. 1(a)).

**FIG. 1.**
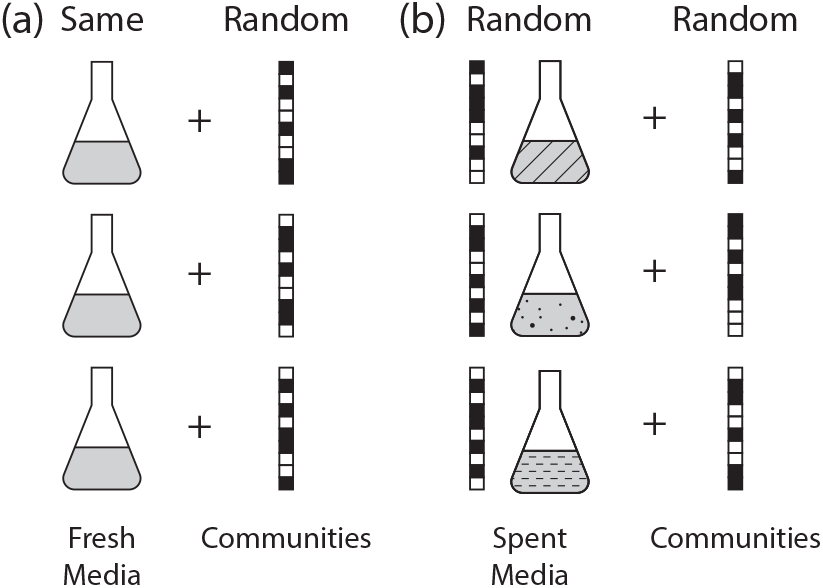
Experimental schemes for (a) the existing protocol of assaying random subsets of the species pool on the same fresh medium, and (b) our proposed random community-media pairing protocol, in which random communities are paired with random spent media. The binary vectors indicate species presence or absence in communities used to generate or pair with random spent media. The second protocol is expected to generate datasets with richer mechanistic information.

However, in biology our goal is often not just to predict (e.g., invasion resistance, or the rate of a metabolic process), but to understand something about the *mechanism* of its collective emergence in the community. While machine learning is increasingly seen as a powerful method to achieve the former, the latter—disentangling the mechanism—remains distinctly more laborious. Here, we ask whether simple machine learning methods, such as regression, can be leveraged to learn mechanistic information.

In particular, this paper focuses on the mechanistic mapping between species and pathway steps. In microbial communities, complex biochemical pathways, such as the degradation of toxic pollutants, are often encoded and executed at the community level [2, 3]. Knowledge about each species’ contribution to these pathways is highly useful for microbial engineering. While such mechanistic information is typically obtained through laborious bottom-up approaches involving extensive metagenomics and metabolite analyses [2, 4–6], here we explore the plausibility of using regression-based, randomized methods to learn mechanistic mappings.

Specifically, we test this idea on cross-feeding relationships. Cross-feeding is a common mechanism underlying the execution of complex biochemical pathways in microbial communities, such as in bisphenol A (BPA) degradation [2]. Spent media analysis is a traditional method for inferring cross-feeding relationships. Spent media refers to the media left after removing the microbes cultivated on it. Comparing the growth of taxa on fresh and spent media yields information about metabolites secreted by some taxa or required by others. For example, many studies use spent media analysis to assess pairwise interactions between taxa [7–10]. Building on this tradition, we combine spent media analysis with the efficiency of randomized sampling, and propose an approach where we pair randomly assembled spent media with random communities (FIG. 1(b)). In this paper, we refer to this approach as the random community-media pairing protocol.

Using a model-based analysis, we show that the random community–media pairing protocol facilitates learning mechanistic information about a collective microbial function. It enables straightforward extraction of a crossfeeding relationship using a simple LASSO-based classifier and successfully identifies the species essential to the collective function. This analysis provides a proof of principle that a regression-like approach can efficiently extract mechanistic information from a complex collective function. In mechanistic microbial study, interpretable algorithms and active learning are two common ways to incorporate machine learning. Our analysis shows that engaging the experimental protocol with machine learning provides a useful additional approach for integration. We further situate this paper in the various strategies of integrating machine learning into mechanistic microbial study, and argue that using ML-synergistic experimental protocols can be a third strategy, in addition to using interpretable ML algorithms and active learning.

## II. *RESULTS*

### A. A Two-step Pathway Component Model

To test the utility of random pairing protocol, we built a model to generate synthetic data. There are two requirements for the model. First, there should be at least two qualitatively distinct ways for a species to contribute to the function, so we can test the ability of our protocol to distinguish between them. Second, the synthetic function should be collective, in the following sense: Define a *functional community* as one for which the value of the functional readout exceeds some threshold (e.g., zero). Define a *minimal* functional community as one where all members are required: dropping any member makes the community non-functional. Now, let *K* denote the number of species in a typical minimal functional community (which need not be unique, due to functional redundancy between species). This *K* quantifies the collective nature of the function. If *K* is low (*K* = 1-2), then examining species individually or pairwise is sufficient to fully understand which species are contributing. The challenge arises when we are unable to reproduce some function in a low-complexity community, as is the case, e.g., for pathogen colonization resistance [11] and efficacious fecal microbiota transplantation [12]. This is the scenario where the traditional approaches struggle, and where randomized methods can be useful [1]. Thus, to test the random pairing protocol, our goal is to select a minimal model that satisfies these two requirements simultaneously.

To satisfy these two conditions, we adopted a model where a complex compound is synthesized through a twostep pathway, each requiring multiple species to run. Examples of such pathways include sugar fermentation in the gut microbiome [13] and BPA degradation [2]. Conceptually, we consider a scenario where a complex compound (“final compound”) is synthesized from another complex compound (“precursor”), as shown in FIG. 2. In our model, both of these steps require cooperation of multiple species, but only the final compound concentration is measurable; the precursor is assumed to be unknown and not measurable. Our goal is to identify which species are participating in the precursor synthesis, and which species participate in the conversion of the precursor into the final compound.

**FIG. 2.**
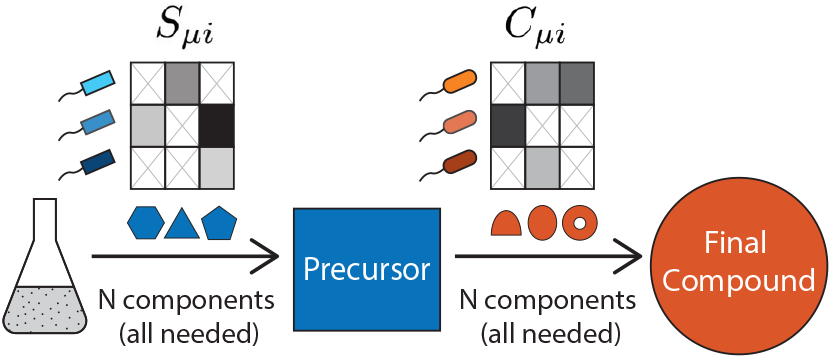
Two-step pathway component model as a minimal computational model with a clear mechanism to be recovered. Precursor is needed for the production of the final compound. For simplicity, species are divided into precursor makers (blue) and precursor converters (orange). The production of precursor and final compound each requires the presence of the complete set of *N* components of the respective pathway (for simplicity, *N* = 10 in both cases). Each species can contribute to multiple components. *S*_*µi*_ and *C*_*µi*_ encode species’ contributions to precursor synthesis and precursor conversion, respectively. Synthetic data generated with this model were used to test the ability of regression to distinguish precursor makers from precursor converters.

We assume that precursor synthesis requires *N* pathway “components.” Each species *µ* may contribute some amounts of components *i* ∈ 1 … *N* of this precursor synthesis pathway, as characterized by a sparse matrix *S*_*µi*_. Successful synthesis requires all *N* components to be present in the community.

Specifically, in the scenario of random subset protocol illustrated in FIG. 1(a), if *x*_*µ*_ ∈ {0, 1 } is the binary vector identifying which species were selected to be part of the community, we postulate the final precursor concentration to be:

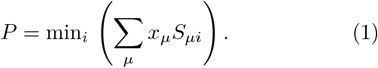

In other words, synthesis is limited by the component with the lowest representation in the community.

Precursor conversion into final compound is implemented in a similar way: each species *µ* contributes components of a precursor conversion pathway, as described by a sparse matrix *C*_*µi*_. The number of components of this pathway is also taken to be *N* for simplicity. The final compound concentration is given by:

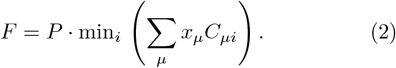

Here, *P* is the precursor concentration computed for this community as above. The final compound concentration *F* is proportional to the amount of precursor, and successful conversion requires all conversion pathway components to be present in the community. Eqs. (1), (2) together described the two-step pathway component model for the random subset protocol scenario in FIG. 1.

For the two-stage random community-media pairing protocol illustrated in FIG. 1, we use a similar set of equations. Specifically, we first create a library of spent media *α* (each defined by binary presence/absence vector of species, which we call 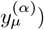, and then perform measurements of the final compound *F* created by a random subset of species 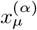 when placed in that spent medium 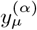. Any precursor and final compound are inherited through the medium:

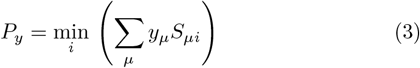

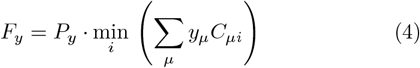

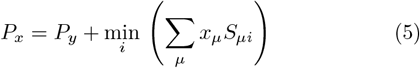

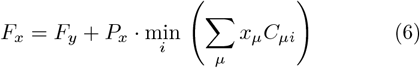

Eqs. (3), (4) describe the synthesis of precursor and final compound in the medium-constructing community, and are identical to Eqs. (1), (2). Eq. (5) says that the precursor created by *x*_*µ*_ adds to the precursor already in the medium (created by *y*_*µ*_). We assume that the precursor-to-final-compound conversion rate is very small (*F* ≪ *P*), so that the conversion to final compound does not appreciably reduce the precursor concentration. This was done for simplicity, and relaxing this assumption does not change the results. Finally, Eq. (6) describes the conversion of precursor into final compound by community *x*_*µ*_, with the new synthesis being proportional to the full precursor concentration in the community (which includes any precursor inherited through the medium).

Note that in this model, we do not track the dynamics of species abundances. Instead, this is directly a model of a complex “functional landscape,” defined as a mapping from either community composition *x*_*µ*_, as in the random subset protocol, or a pair of community composition and spent media label (*x*_*µ*_, *y*_*µ*_), as in the random community-media pairing protocol, into a functional property (in this case, final compound concentration *F*) [1, 14]. The mapping we defined is highly non-linear due to the minimum function, and since the matrices *S* and *C* are sparse, the production of precursor and the conversion to final compound usually each requires multiple species to be present simultaneously (higher-order interactions). The precursor synthesis matrix *S* and precursor conversion matrix *C* are the mechanistic “ground truth” that characterizes how a given species is contributing to the community-level function *F* (final compound concentration).

For simplicity, here we assume that each species may contribute either precursor synthesis components or precursor conversion components, but not both. (If row *µ* of matrix *C* has a non-zero entry, the corresponding row in matrix *S* is all zeros, and vice versa. The zero half of *S*_*µi*_ corresponding to precursor converters and the zero half of *C*_*µi*_ corresponding to precursor producers are omitted in FIG. 2 for simplicity.) This assumption turns the problem of distinguishing species’ contribution types into a binary classification problem, which allows for an easier definition of a performance metric. Specifically, we evaluate performance using the Matthews Correlation Coefficient (MCC), calculated between the true classification of species as a precursor-maker or a precursor converter and the algorithm’s best guess. The MCC is a standard method of evaluating the performance of binary classifiers [15].

### B. Random Subset Protocol Is Insufficient for Mechanistic Species Classification

Our model implements a minimal scenario where being able to predict *F* does not, on its own, answer the mechanistic question of how each species contributes to the function. Indeed, if we observe that adding a species *µ* to some community increases the final compound concentration *F*, this could be for either of two reasons: *µ* might contribute to precursor synthesis, or might improve its conversion into final compound. In this section, we show that using random subset protocol poses a challenge to distinguish between the two types of contributions.

We first check that our pathway component model is an example of a scenario where linear regression succeeds at prediction. We define richness as the typical number of species present in a randomly assembled community. We also introduce density, which refers to the probability that a precursor producer is capable of contributing to a given precursor-production component (so named because it relates to the fraction of nonzero entries in the pathway matrix). For simplicity, we use the same probability for a precursor converter contributing to a precursor-conversion component. A density of 1 corresponds to a fully redundant system in which every precursor maker can independently produce the precursor, and every precursor converter can independently carry out the conversion. As density decreases, production of the final compound becomes increasingly dependent on collective contributions from multiple species. We ran 500 trials for each experiment and performed LASSO-regularized linear regression with 10-fold cross validation [16, 17] over 450 trials and tested prediction performance over the rest 50 trials. We repeat this process 500 times for each pair of richness and density. FIG. 3(a) reports the mean of the prediction performance (measured in terms of the Pearson correlation coefficient between the predicted *F* and the measured *F*). The panel shows that LASSO regression succeeds at prediction over a wide range of parameter space (the decrease in performance in the lower-left region is expected, as communities sampled are often non-functional). Hence, the model is within the desired scenario where regression is capable of prediction.

**FIG. 3.**
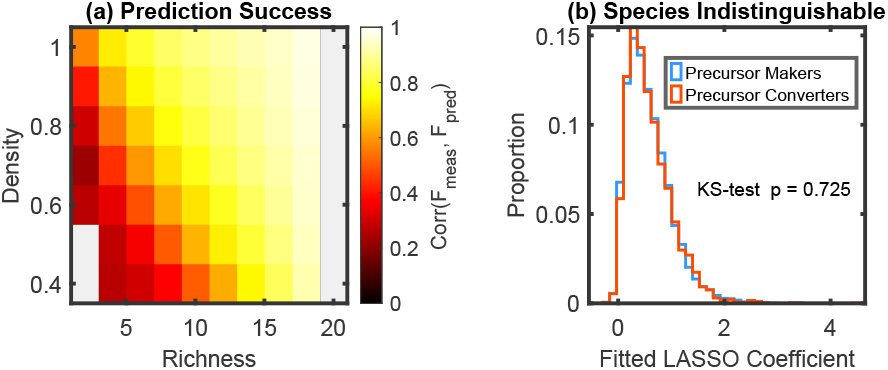
Sufficient for functional prediction, the existing protocol is insufficient for mechanistic species classification. (a) Simple regression is sufficient for function prediction. Despite the complexity of collective microbial functions, previous research shows that simple regression over presenceabsence information of random communities and their respective functional readouts is sufficient for function prediction. As shown, our model operates in the parameter regime where simple LASSO regression succeeds at function prediction, consistent with prior empirical studies. (Density controls the probability of a species being able to produce a component. Richness refers to the typical number of species drawn from the species pool into a community. 20 species in the pool. 500 trials for each dataset, 500 datasets for each parameter pair. Dataset training-testing split is 90-10.) (b) Despite the success in prediction, random community data does not readily provide mechanistic information about how species contribute to the function. Statistically, the distributions of the fitted coefficients for different types of contributors are indistinguishable (KS-test: p=0.725; see FIG. S1 for more analysis). This indistinguishability persists even when using more advanced algorithms and incorporating additional prior information (see Materials and Methods and FIG. S2, S3).

However, for random subset datasets, while simple LASSO regression already succeeds at prediction, such datasets are not very informative about how species contribute to the final compound concentration *F*. Indeed, the distributions of the LASSO-fitted coefficients for the two types of species are statistically indistinguishable (see FIG. 3(b), S1). This is still the case if we switch to a more advanced algorithm based on random forest regression (FIG. S2). Even when we recast the problem as a supervised learning task, providing the model with prior knowledge of half of the species’ identities, it still fails to classify the remaining unknown species (FIG. S3). We therefore ask whether a different experimental design could facilitate the extraction of mechanistic information.

### C. Random Community-media Pairing Succeeds at Extracting Mechanistic Information

The alternative experimental design we propose is the random community-media pairing protocol, as illustrated in FIG. 1(b). We generated synthetic datasets with the new protocol, again based on the two-step pathway component model. Each data point in such new datasets consists of two binary vectors,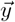 and 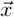 and a scalar *F*, the level of the final compound at the end of each trial. As mentioned before, we assume that *F* is the only quantity measured in the experiment.

We first checked the behaviors of LASSO-regression over such datasets. Over each dataset (equivalent to one set of experiment), we trained LASSO regression with 10-fold cross validation, and LASSO regression returned two coefficients for each species, *α* and *β*, with *α* corresponding to 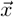 and *β* corresponding to 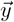. As FIG. 4(a) shows, (*α, β*)’s tend to split on the two sides of the 1-1 line, with the splitting highly correlated with the types of the corresponding species. Inspired by this observation, we configured a classifier: for each species *µ*, if *β*_*µ*_ *> α*_*µ*_, then *µ* is classified as a precursor maker; conversely, if *α*_*µ*_ *> β*_*µ*_, then *µ* is classified as a precursor converter. We then checked the performance of this simple LASSO-based classifier in extracting mechanistic information about the collective function. If *β*_*µ*_ = *α*_*µ*_, we label a species as “unclassified” and treat this as a classification error for conservative evaluation of classification performance.

**FIG. 4.**
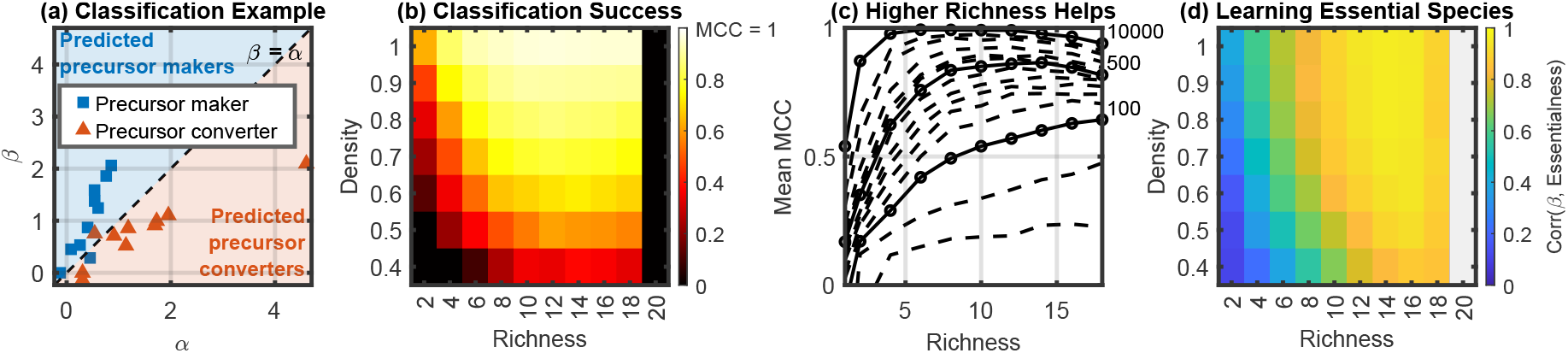
Random community-media pairing succeeds at extracting mechanistic information. (a) Illustration of classification process using LASSO-fitted coefficients over random community-media pairing datasets. Each species has two fitted coefficients: *α* and *β*, corresponding to the random communities stage and the random media stage, respectively. If *β > α*, the species is classified as a precursor maker; conversely, if *α > β*, the species is classified as a precursor converter. Rest of this figure applies this classifier. (b) With random community-media pairing, mechanistic information was easily extracted. LASSO-based classification outputs were compared with ground truth, performance measured with Matthews Correlation Coefficient (MCC). 500 trials for each dataset, 500 datasets for each parameter pair. Plotted are means over the 500 datasets. Low performance at the lower-left corner is expected, as the assembled communities are usually non-functional (FIG. S4). Simpleness of this LASSO-based classifier highlights the mechanism-revealing strength of random community-media pairing. (c) Including more species in communities is helpful. Shown are performance curves with different trial numbers for density = 0.8. Bold curves correspond to trial number = [100, 500, 10000]. Data points covers richness = [1, 2, 4, 6, 8, 10, 12, 14, 16, 18]. From bottom to top, curves shown correspond to trial number = [30, 50, 100, 150, 200, 250, 300, 400, 500, 600, 700, 1000, 2000, 3000, 10000]. Increasing community richness significantly increased classification performance, especially for low-trial-number scenarios. (d) Random community-media pairing helps extract additional mechanistic information about essential species. Species essentialness is defined with the likelihood of each component being the bottleneck that limits the production of either the precursor or final compound, along with species’ abilities to produce the component (see Materials and Methods). As shown, for predicted precursor makers, their essentialness values correlate strongly with their fitted *β* values. The same holds true for predicted precursor converters (showing a high correlation with fitted *α*’s), and both correlations are statistically significant (see FIG. S6). Such high correlations mean that random community-media pairing enables further mechanistic understanding of the species pools.

As shown in FIG. 4(b), the classification has a wide success range in the density-richness parameter space. Here, richness refers to the typical number of species used to construct a spent medium and the typical number of species in a subset community (same richness for simplicity). The sub-optimal performance at the lower-left corner is understandable, as most communities assembled in this parameter regime are non-functional (FIG. S4). As expected, classification performance generally decreases with density. Lower density corresponds to higher collectivity of function, as typically more species are needed to form a functional community (see FIG. S5 for details). However, this performance drop can be compensated by increasing community richness (which yields more highfunctioning examples). That the LASSO-based classifier continues to succeed at high-collectivity conditions highlights its capability to extract mechanistic information about collective microbial function.

Note that FIG. 4(b) demonstrates that performance loss in low-density (high-collectivity) regime can be compensated by increasing the richness of trial communities. FIG. 4(c) further highlights the general increase of classification performance with richness. In other words, the protocol presented here succeeds not *despite* high richness of trial communities, but rather *because* of it, as richness generally improves performance. (The slight nonmonotonicity at high richness observed for very large trial numbers is expected: for any finite species pool, once the richness reaches half of the total number of species, increasing it further will make the assembled communities increasingly alike; for instance, there is only one possible community when richness equals the total number of species.) Although our minimal model is not meant as a faithful representation of any particular microbial system, these observations indicate the promise of randomized sampling of large subsets of species pool in the mechanistic study of collective microbial functions.

In addition to classifying species, the LASSO-based classifier over data generated by random community-media pairing also extracts mechanistic information about the “essentialness” of each species for the collective function. In our pathway component model, the production of precursor is limited by the least available component in the system; the same holds true for the conversion from precursor to final compound. We call such components bottleneck components. To assess whether the classifier can learn about species essential to the collective function, we quantify the “essentialness” of a species as the sum of the species’ ability to generate each component, weighted by the likelihood of that component being the bottleneck component (see Materials and Methods for details). The two sets of statistics produced by the classifier, *β*’s and *α*’s, are found to strongly correlate with the essentialness of predicted precursor producers and predicted precursor converters, respectively (FIG. 4(d), S6). It is noteworthy that, for high richness, the two statistics remain highly correlated with species essentialness even when the classification performance is sub-optimal (e.g., density = 0.4) (FIG. 4(b, d), S6). Thus, at least in our model, the protocol succeeds at identifying species essential to the collective function.

## III. DISCUSSION

Together, these results provide a proof of principle that the merit of regression-like methods for studying collective microbial functions may be extended beyond approximate predictions, and towards extracting mechanistic information. Specifically, we show *in silico* that a regression-like method over randomized data can reveal mechanistic insights about a collective microbial function. We built a minimal two-step pathway model that generated synthetic data following the random pairing protocol, which randomly pairs spent media with species subsets. We showed that LASSO regression over this dataset is enough to classify species by the way of their contribution to the collective function—either contributing to the generation of the precursor or helping to convert the precursor to the final compound. The LASSO-fitted coefficients also extract additional mechanistic information about essential species that are most likely to cause bottlenecks in either the production of the precursor or the production of the final compound. These results demonstrate how a shift in experimental design— from random subset protocol to random community-media pairing—can enable the extraction of mechanistic information.

In biological studies seeking mechanistic insight, machine learning has been applied in two principal ways. The first one is to develop algorithms whose output would be more mechanistically interpretable [18–20] (FIG. 5(a)). The second approach, known as active learning [21–23] (FIG. 5(b)), is to iteratively use machine learning to predict which additional data points are the most informative and hence should be measured next. Complementing these approaches, our results highlight the potential of integrating machine learning at the level of a synergistic experiment design (FIG. 5(c)). Specifically, in our analysis, even an extremely simple algorithm (linear regression) can reveal much more insight with an appropriately modified data collection protocol.

**FIG. 5.**
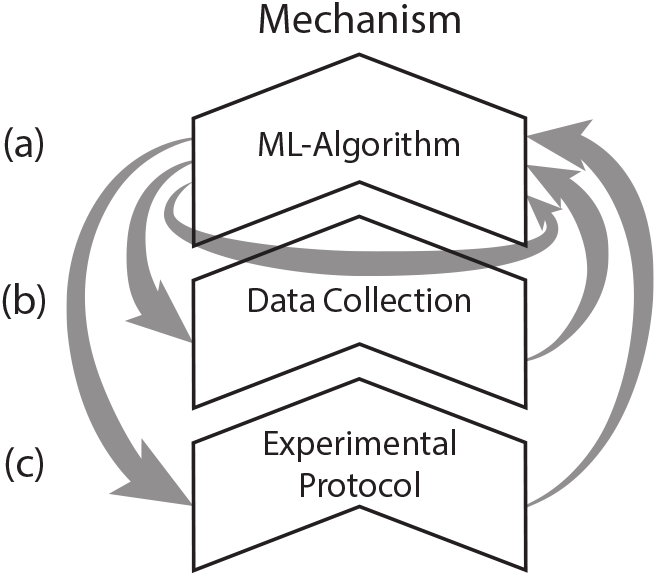
Tiers of machine learning (ML) integration into studies of collective microbial function. The dominant two approaches of harnessing ML approaches for deriving mechanistic bioloigcal insight are (a) improving the interpretability of ML algorithms and (b) using ML to guide data collection through active learning, which predicts the most informative measurements to measure next. (c) Our results highlight a complementary third strategy: “ML-synergistic experimental design.” In the example we presented, the two-step experimental design allows even simple regression models to recover mechanistic information about the collective function.

In real-world experiments, there are two easily available improvements for extracting mechanistic information. First, after assembling a spent medium, it is natural to measure its composition. How this spent medium differs from the fresh medium yields additional information about the species being used to make the spent medium. Another improvement is that one can use an algorithm more advanced than regression. In this paper, we were less concerned by efficiency of inference, but intentionally limited our inference to the minimal “one functional measurement per pairing” and simple LASSO regression to illustrate the core idea.

## IV. MATERIALS AND METHODS

### A. Simulation details

We worked with microbial pools of 20 species, consisting of 10 precursor producers and 10 precursor converters. Each pool is defined by a pair of pathway–component matrices, *S* and *C*. Density controls the percentage of non-zero entries in *S* and *C*. The value of each nonzero entry is drawn from an exponential distribution with mean 1. A new pair of *S* and *C* matrices is generated for each dataset.

All simulations were performed in MATLAB (Mathworks, Inc.). The associated code, simulation data and scripts to reproduce all figures in this work are publicly available [24].

### B. Matthews Correlation Coefficient

This paper uses Matthews Correlation Coefficient (MCC) to evaluate classification performances. MCC is a robust single-value classification metric for balanced datasets, in which positive and negative cases hold equal importance [15]. MCC can be expressed as:

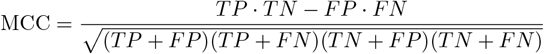

where *TP, TN, FP*, and *FN* correspond to true positives, true negatives, false positives, and false negatives, respectively. MCC = − 1 means perfect classification; MCC = −1 means perfect mis-classification; and MCC = 0 corresponds to random classification. We arbitrarily regarded being precursor makers as the positive cases, and being precursor converters the negative. This choice has no impact on the evaluation, as MCC is symmetrical with respect to positive and negative cases. Additionally, we treated non-classification as false classification for conservative evaluations of MCC’s.

### C. Random Forest Regression classification attempts with random subset datasets

We also used classifiers based on random forest regression over the random subset datasets.

#### Unsupervised

Given the classification failure of random forest regression on random subset dataset, we further checked if a more sophisticated unsupervised algorithm using random forest regression (RFR) will be able to succeed. We first trained RFR for function prediction (performance shown in FIG. S2(a)). Then, to leverage the intuition that precursor makers and precursor converters contribute to the collective function differently, we used the trained RFR model to construct species contribution profile vectors, which we define as the RFR-predicted effect of perturbing presence/absence of the respective species across all trials. K-means over species contribution profile vectors then divided species into two clusters. Each cluster was leniently labeled with the true majority identity of its constituents; species were classified with cluster labels. FIG. S2(b) records the consistent failure of this method at classification.

#### Supervised

We further lessen the difficulty of the task by allowing prior knowledge about the identities of half of the species pool (5 known precursor makers and 5 known precursor converters) and turning the task into a supervised classification problem. Same method as aforementioned was used to generate the contribution profile vectors. Linear SVM and logistic regression were trained on the contribution profile vectors of the 10 known species and then used to classify the 10 unknown species. Classification performance was then evaluated with regard to the 10 unknown species. FIG. S3 shows the continued failure of classification.

These failures with more advanced classifiers and additional information demonstrate the underdesirability of random community protocol for extracting mechanistic information.

### D. Bottleneck components and species essentialness

Bottleneck components refer to the components *i* that limit either the production of precursor or the conversion of precursor to the final compound. In other words, bottleneck components are the components at which the minimums in Eqs. 3–6 are taken. These bottleneck events are recorded separately for the random media phase and the random community phase and then pooled together. A small Gaussian noise (*ε* = 10^−10^) is intentionally added when determining the bottleneck component in order to break potential degeneracies when two or more components have identical amounts available; without this perturbation,`min()` in MATLAB would always return the earliest component in the list. This noise is used only for selecting the bottleneck index and is not included in the calculation of precursor production or conversion to the final compound. Additionally, a bottleneck component on the conversion step is only returned when precursor is present in the system.

For a component *i* in the precursor synthesis matrix *S*, its bottleneck likelihood *b*_*S*_(*i*) is defined as the number of times component *i* was identified as a bottleneck, normalized by the total number of bottleneck events observed for that pathway step. *b*_*C*_(*i*) is defined analogously for components in the precursor conversion matrix *C*. For each pair of pathway matrices *S* and *C*, we simulate 10,000 additional trials to estimate these likelihoods, so that the estimates are independent from the trials used to train the LASSO regression.

Species essentialness is then defined using these bottleneck likelihoods. The essentialness of species *µ* is defined as

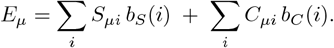

Thus, a species is regarded as more essential if it contributes strongly to components that are frequently limiting in the system.

## ACKNOWLEDGMENTS

We thank L. Graham and J. Moran for helpful discussions. This work was supported in part by NSF grant PHY-2340791. S.K. acknowledges the National Institute of General Medical Sciences R01GM151538 and support from the National Science Foundation through the Center for Living Systems (grant no. 2317138). S.K. and M.T. also acknowledge support from ARO W911NF2510213.

## SUPPLEMENTAL MATERIAL

This Supplemental Material contains additional figures referenced in the main text. Figures are labeled Fig. S1, Fig. S2, etc.

**FIG. S1.**
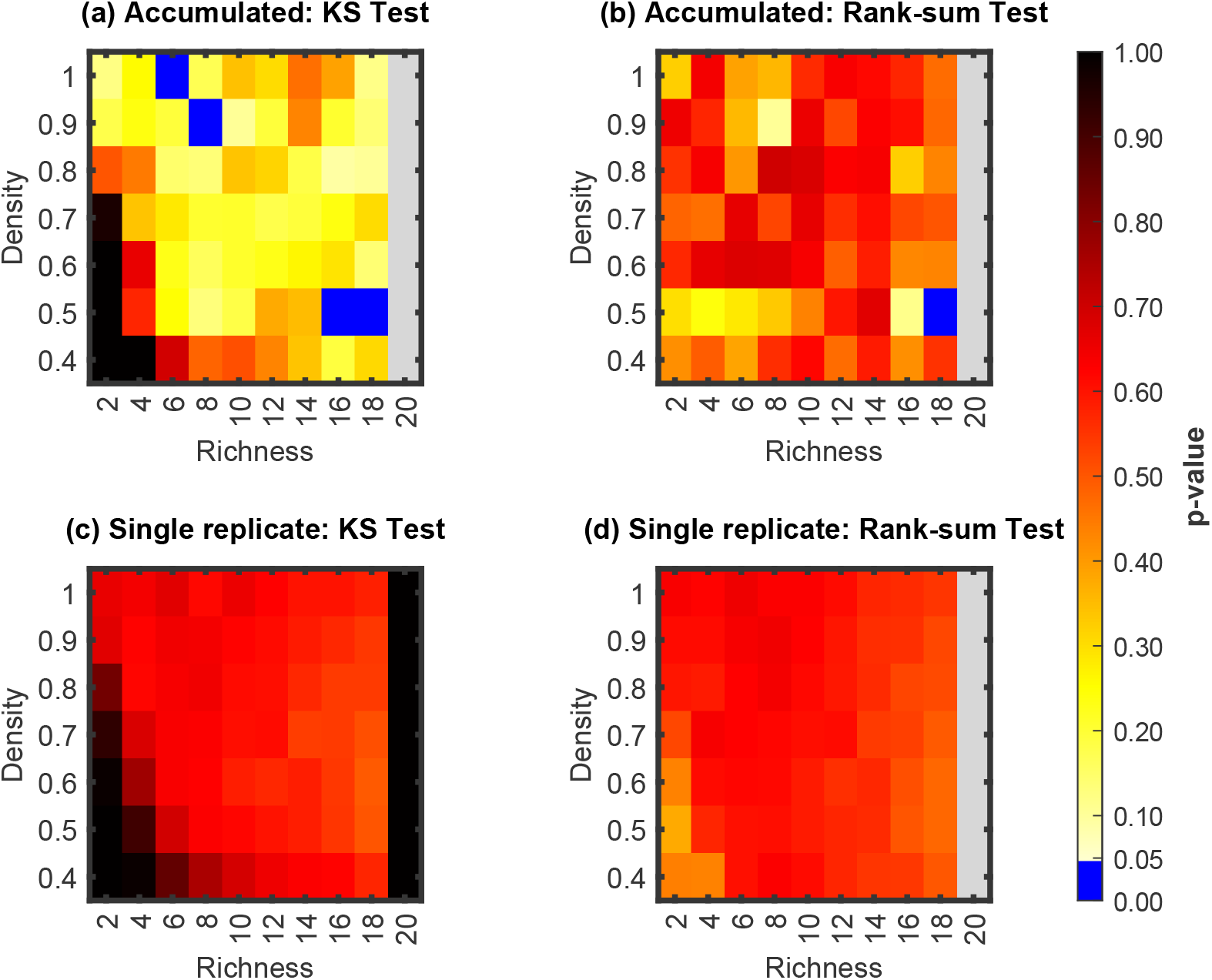
For random-subset datasets, fitted LASSO coefficients offer little statistical power for distinguishing precursor producers from precursor converters. (a) When LASSO-fitted coefficients from all datasets at a given richness–density pair are pooled together, the distributions for the two species types are only rarely statistically distinguishable (Kolmogorov–Smirnov test, or KS test for short). (b) Moreover, one distribution only rarely tends to take systematically larger values than the other (Wilcoxon rank-sum test, or rank-sum test for short). (c) At the level of individual datasets, which is more directly relevant experimentally, the two distributions again appear statistically indistinguishable under the KS test, and (d) neither distribution tends to be systematically larger under the rank-sum test. Together, these results indicate that fitted LASSO coefficients cannot meaningfully separate the two biological classes using any simple cutoff. For each richness–density condition, 500 independent datasets were simulated. In panels (a) and (b), we performed 500 bootstrap resamplings at each parameter pair; each resampling draws 500 datasets with replacement, pools the corresponding LASSO coefficients across all species, and computes KS and rank-sum *p*-values. The median *p*-values across bootstrap replicates are shown. Panels (c) and (d) instead compute *p*-values within each individual dataset, comparing precursor vs. converter coefficients; the plotted values are the mean *p*-values across the 500 datasets at each parameter pair.

**FIG. S2.**
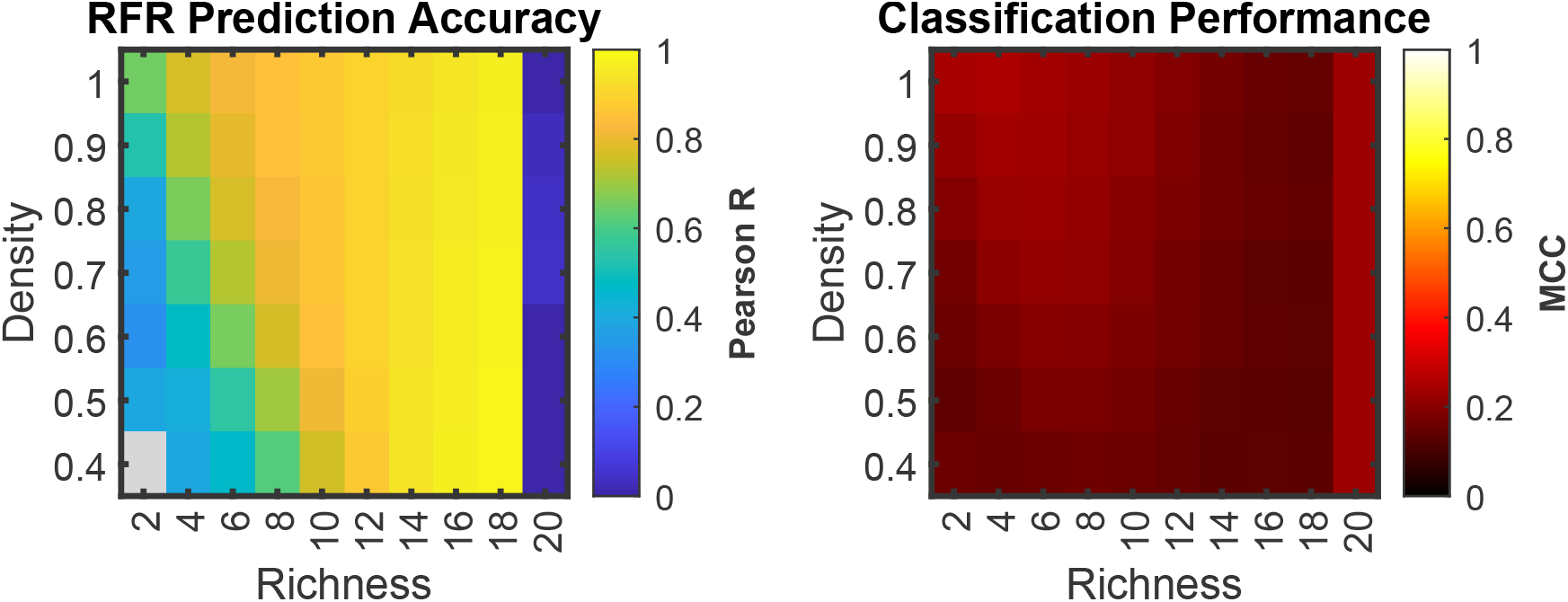
Prediction success of random forest regression (RFR) over random subset dataset and classification failure of unsupervised clustering of corresponding RFR-generated contribution profile vectors. At richness = 20, the clustering yields MCC = 0.23, corresponding to singling out one species, with all the other 19 species grouped in another group. This behavior arises from the imposed requirement of k-means clustering to partition the data into exactly two clusters (*k* = 2).

**FIG. S3.**
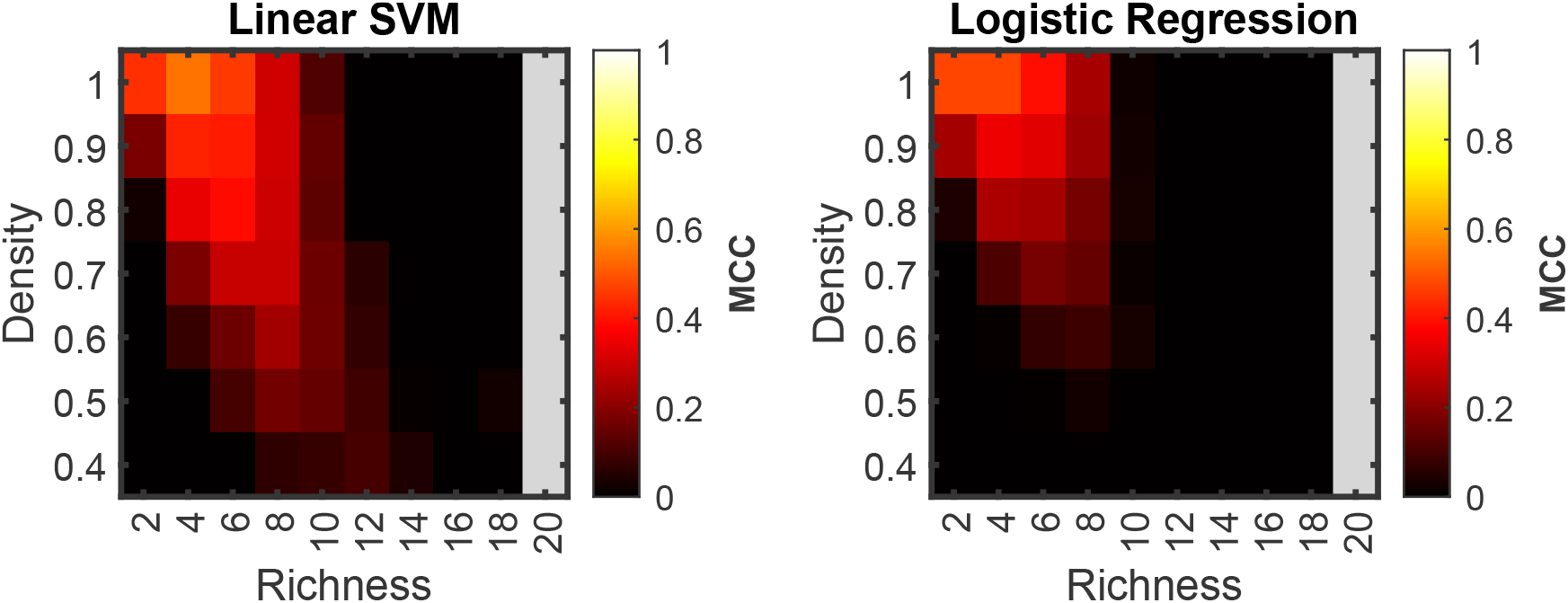
Classification performance of linear SVM and logistic regression on contribution profile vectors predicted by random forest regression.

**FIG. S4.**
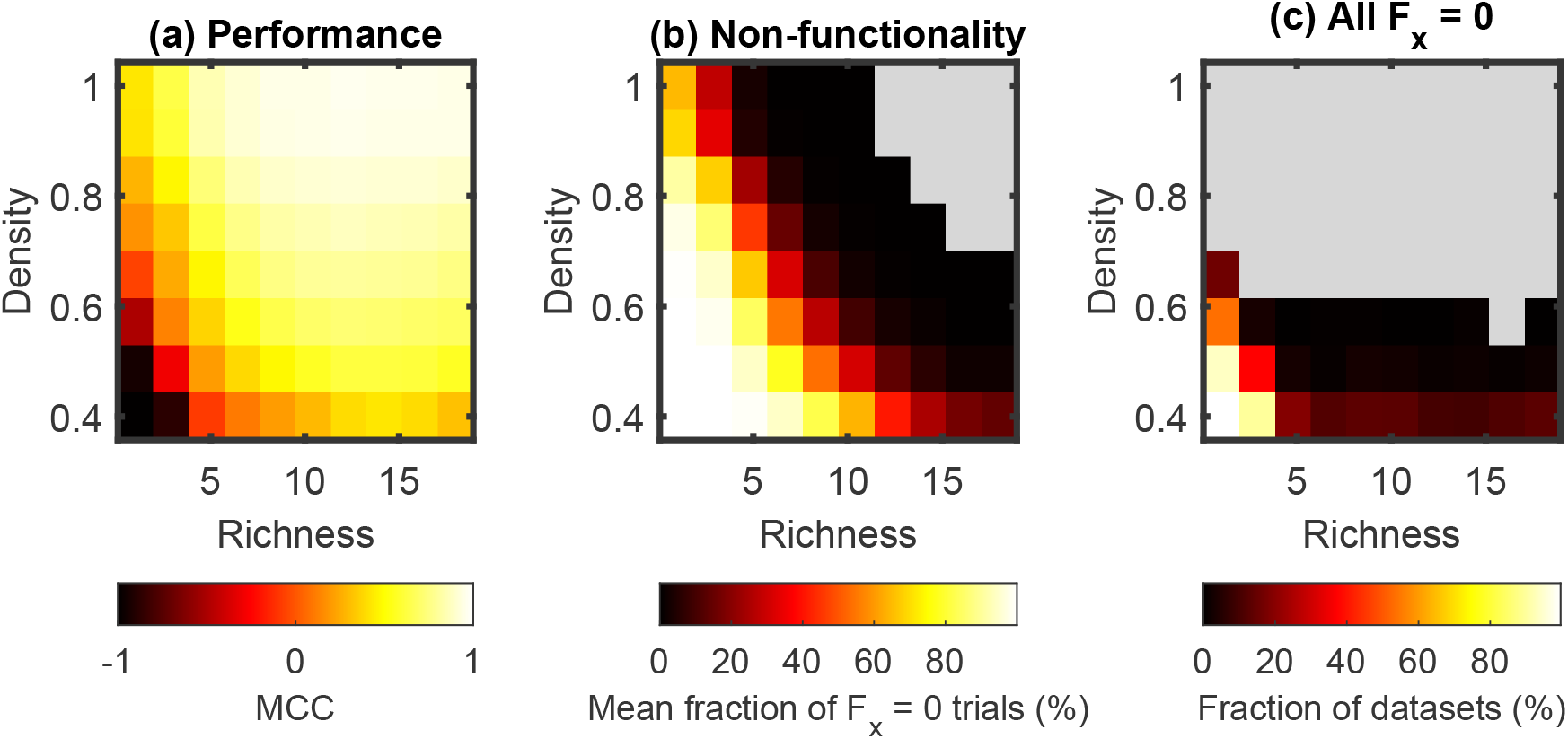
Low classification performance at the low-density-low-richness regime is expected due to increased non-functionality of assembled communities. (a) Reproduction of Fig. 4(b) with colormap range over [-1,1], instead of [0,1]. (b) Average percentage of non-functional trials (defined as *F*_*x*_ = 0, meaning that both *µ*_*x*_ and *µ*_*y*_ are non-functional communities). As shown, as density and richness decrease, communities assembled are increasingly likely to be non-functional. This coincides with the gradual decline in classification performance. (c) Percentage of datasets whose 500 trials are all non-functional. For a dataset, when all 500 *F*_*x*_’s are zero’s, LASSO regression will fail and produce zeros for all fitted coefficients. This leads to all species being non-classified and counted as classification errors, resulting in MCC = − 1. This explains the sharp drop in classification performance at the lower-left corner.

**FIG. S5.**
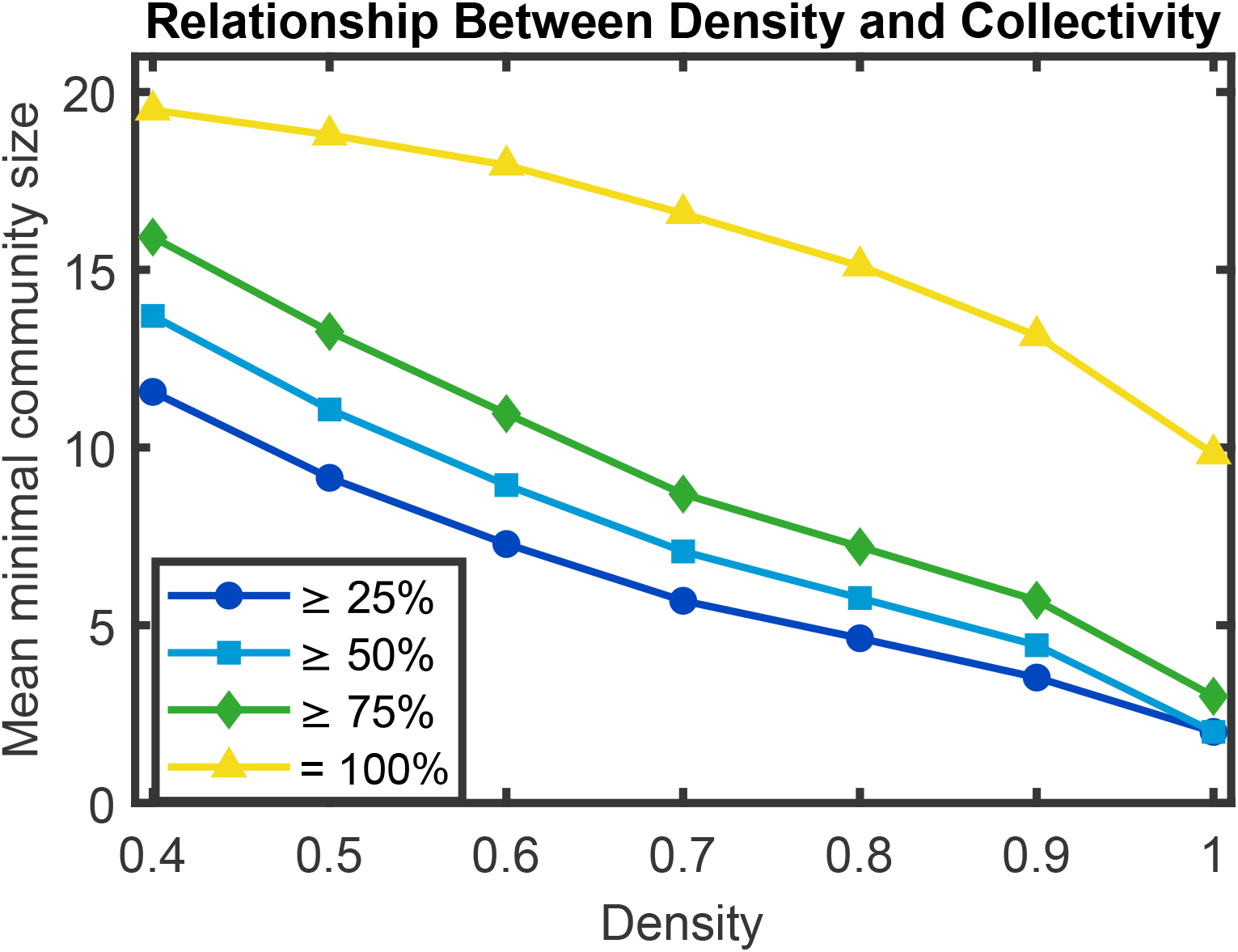
Lower pathway density corresponds to a more collective regime, in which larger communities are typically required to produce the final compound. For each density, we generated 500 independent pairs of pathway matrices (*S, C*) and, for each pair, evaluated the fraction of 10,000 randomly assembled communities of size *k* (with *k* = 1, …, 20 species) that produced the final compound. Plotted are the mean minimal community sizes required to achieve at least 25%, 50%, 75%, or 100% probability of producing the final compound.

**FIG. S6.**
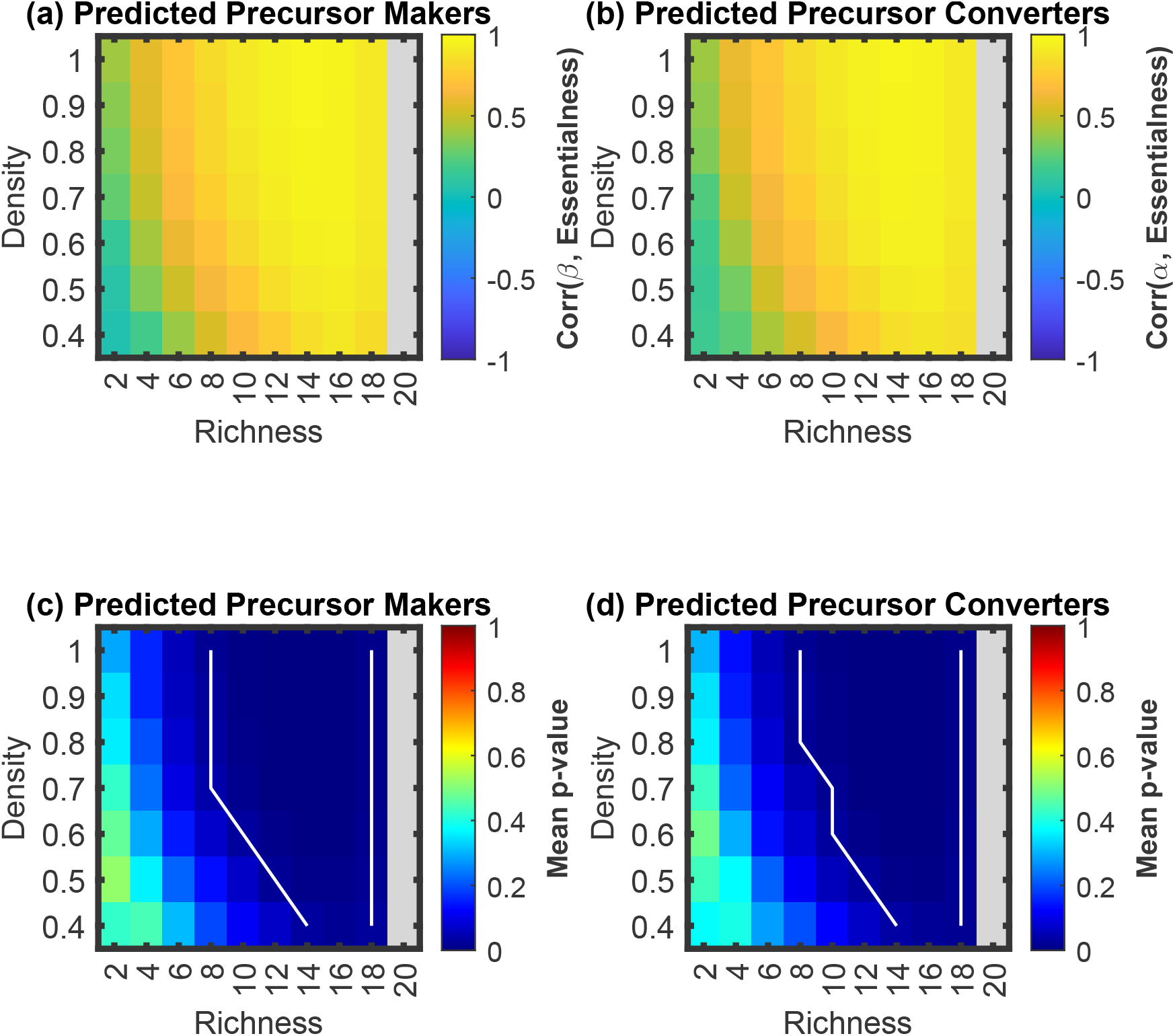
LASSO regression applied to random community–media pairing datasets recovers additional mechanistic information about species essentialness. (a) Among predicted precursor makers, the LASSO-fitted *β*’s correlate strongly with species essentialness across a broad range of richness–density conditions. (b) Similarly, among predicted precursor converters, the LASSO-fitted *α*’s show strong correlations with essentialness over a wide parameter regime. (c,d) These correlations are statistically significant, as indicated by the white contour marking the *p* ≤ 0.05 significance boundary. All quantities shown are averages over 500 replicates.

